# Insectivores exhibit superior microbial transmission efficiency and elevated zoonotic risk by 2035 relative to rodents and bats

**DOI:** 10.1101/2025.09.10.675328

**Authors:** Hongfeng Li, Zheng Y.X. Huang, Jie Lan, Li Hu, Zhilin Wang, Yingying Wang, Luca Santini, Daniel J. Becker, Fuwen Wei, Yifei Xu

## Abstract

Wild small mammals represent critical sources of zoonotic infections due to their high diversity, global distribution, and proximity to humans. Nevertheless, significant knowledge gaps persist in characterizing pan-taxonomic microbial richness and sharing dynamics, particularly regarding ecologically critical yet understudied Eulipotyphla (true insectivores). Here we take a macroecological approach to compare how microbial hosting and transmission differ across insectivores, rodents, and bats, and what ecological factors drive such disparities. We find that insectivores host comparable microbial richness to rodents and bats while exhibiting superior intra- and cross-order transmission efficiency. Urban adaptation, geographic range area, and longevity are shared drivers of microbial richness and transmission across these host orders, while greater body mass and shorter gestation time specifically are positive predictors of these outcomes within insectivores. Climate change projections identify insectivores as primary transmission hosts in new high-latitude hotspots by 2035, including parts of the US, Canada, and Russia, posing greater zoonotic threats than rodents or bats. Our findings challenge the prevailing paradigm that prioritizes rodents and bats as special zoonotic reservoirs, establishing insectivores as critical but overlooked players in disease ecology. Collective proactive surveillance of insectivores, rodents, and bats is imperative for forecasting emerging zoonotic threats and informing global risk assessment frameworks.

## Introduction

Most emerging infectious diseases in humans originate through cross-species pathogen transmission from animal hosts^1,2^. Wild small mammals, notably species from the orders Rodentia (rodents), Chiroptera (bats), and Eulipotyphla (“true insectivores”, hereafter insectivores), represent critical sources of zoonotic pathogens due to their high diversity, global distribution, and proximity to and occasional-to-frequent interactions with humans^3–5^. Most work-to-date has focused on the diversity and human infectivity of viruses in particular ^2,6–8^, with relatively less attention to other microbial taxa^9,10^. As such, the diversity and transmission of microbes more broadly within and across wild small mammals remain poorly understood. A better understanding is of central importance for mitigating and preventing future infectious disease outbreaks.

According to the special reservoir hypothesis, rodents and bats are distinct hosts that disproportionately maintain zoonotic pathogens and transmit them to humans^11^. High-profile viruses have been especially well characterized in these host orders and include but are not limited to severe acute respiratory syndrome–related coronaviruses (SARS-related CoVs), henipaviruses, lyssaviruses, and hantaviruses^12–15^. This ability to host such virulent viruses, often without showing disease, likely comes from their special ecological traits, such as high metabolism, torpor, and unique immune systems^16–18^, although the speciose nature of these orders also may explain the high viral diversity^11^. In contrast to these well-studied groups, insectivores have recently been shown to also harbor a number of important zoonotic viruses, such as hantaviruses, Borna disease virus (BoDV), Middle East respiratory syndrome–related coronaviruses (MERS-related CoVs), severe fever with thrombocytopenia syndrome virus (SFTSV), and Langya henipavirus^19–23^. Some insectivore species even host more viruses than rodents and bats^3^. Additionally, among all mammalian orders, insectivores hold the second-most central position for virus sharing, posing a potential threat to public health through spillover and emergence^8^.

While prior investigations have advanced our understanding of viral diversity and transmission in wild small mammals, research effort has been strongly skewed towards rodents, bats, and this particular pathogen group. This has resulted in significant gaps in characterizing pan-taxonomic microbial richness and sharing dynamics, which is especially pronounced for understudied but ecologically critical groups such as insectivores and for the broader (non-viral) microbial community. Specifically, it remains unexplored whether the pattern of microbe hosting and transmission differs across insectivores, rodents, and bats as well as what ecological factors underpin these potential disparities. Filling these knowledge gaps is essential for building predictive frameworks to optimize zoonotic surveillance resource allocation and preempt spillover events. Such predictive efforts are increasingly urgent under climate change, which can alter host distributions, abundance, and contact rates, thereby reshaping the landscape of microbial transmission risk within and between host species^24,25^.

Here, we took a macroecological approach to comprehensively investigate microbe hosting and transmission patterns across insectivores, rodents, and bats. We aimed to (i) compare the patterns of microbe hosting and transmission between these three orders, (ii) dissect ecological factors underlying the differences, and (iii) predict geographic hotspots of microbial transmission risk under several climate change scenarios. Our work highlights wild small mammals, in particular insectivores, that present risks of pathogen spillover to humans and provides a scientific basis for targeted surveillance and early warning systems.

## Methods

### Data collection

We collected host–microbe data for insectivores, rodents, and bats from multiple sources prior to 17 November 2024. Data sources included the Global Virome in One Network (VIRION)^26^, Enhanced Infectious Diseases Database (EID2)^27^, GenBank database, and a specialized insectivore database^8^. All records were merged and deduplicated to ensure data accuracy. Host and microbe species names were standardized to Latin names using the NCBI Taxonomy Database for consistency. Microbes included viruses, bacteria, protozoa, and fungi. The final database comprised 1,424 insectivore host–microbe interactions (922 microbes across 132 host species), 8,937 rodent host–microbe interactions (4,678 microbes across 683 host species), and 9,113 bat host–microbe interactions (4,686 microbes across 526 host species; Fig. 1a). Human-associated microbes were defined as microbes reported in humans in at least one of the VIRION, EID2, or GenBank databases^2,8,28^.

**Figure 1.**
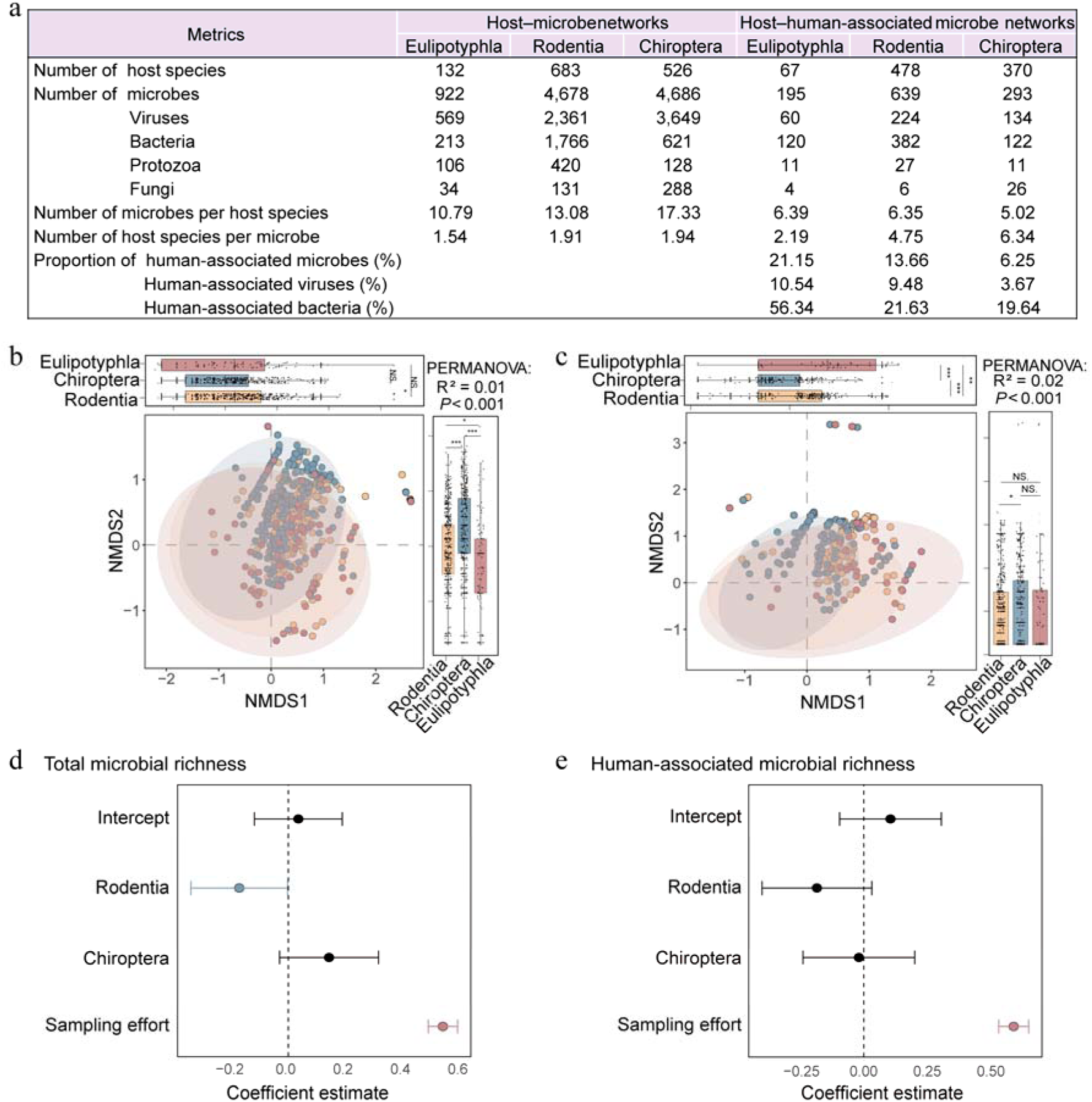
Microbial diversity in insectivores, rodents, and bats. a) Statistical metrics of host–microbe networks. Proportion of human-associated microbes (%) denotes the proportion of human-associated microbes relative to total microbes. NMDS analysis of b) total microbial diversity (insectivores, *n* = 132; rodents, *n* = 683; bats, *n* = 526) and c) human-associated microbial diversity (insectivores, *n* = 67; rodents, *n* = 478; bats, *n* = 370) among three host orders based on Jaccard distance. Ellipses represent 95% confidence intervals (CIs) around group centroids. Boxplots display data distributions with the central line indicating the median, the box boundaries representing the interquartile range (IQR, 25th to 75th percentiles), whiskers extending to the most extreme data points within 1.5×IQR from the box, and dots beyond the whiskers shown as outliers. Asterisks denote statistical significance: ns, *P* > 0.05; *, *P* ≤ 0.05; **, *P* ≤ 0.01; ***, *P* ≤ 0.001. The coefficients estimates and 95%CIs for d) total microbial richness (*n* = 1100) and e) human-associated microbial richness (*n* = 785) derived from the PGLS models. Circles represent coefficient estimates, and lines indicate 95%CIs. Statistical significance: red, *P* ≤ 0.001; yellow, *P* ≤ 0.01; blue, *P* ≤ 0.05; black, *P* > 0.05.

### Microbial diversity across host orders

Variation in host species-level microbial composition across host orders was examined through non-metric multidimensional scaling (NMDS) analyses based on Jaccard distance metrics, implemented with the *vegan* package in R (v4.3.1)^29^. Permutation multivariate analyses of variance (PERMANOVA) were used to test differences in beta diversity. To correct for potential sampling bias, rarefaction curve analysis of microbial richness in our three host orders was conducted based on the number of sampled host species within insectivores, rodents, and bats. We derived a phylogeny of all included host species from the PHYLACINE database^30,31^ and then used phylogenetic generalized least squares (PGLS) models to analyze microbial richness while controlling for evolutionary relatedness. We first estimated phylogenetic signal using Pagel’s λ from an intercept-only PGLS model with the *caper* package^32–34^, where a λ of 0 indicates no phylogenetic signal, while a λ of 1 suggests strong phylogenetic dependence. We then fit PGLS models in the *phylolm* package to analyze microbial richness as a function of host order while controlling for log_(x+1)_-transformed sampling effort^35^. Sampling effort was defined as the number of PubMed citations with the Latin name of each host species.

### Microbe-sharing network

Microbe-sharing networks were constructed to characterize cross-species transmission dynamics across three host orders studied. Bipartite host–microbe networks were independently generated for three host orders collectively and for each order individually, with separate analyses performed for both total and human-associated microbes. Nodes represented host or microbe species, while edges indicated host–microbe associations. These bipartite networks were subsequently projected into unipartite microbe-sharing networks where nodes exclusively represented host species and edge weights quantified the number of shared microbial species between host pairs. Topological characterizations were conducted at network and node levels. Network-level metrics included assortativity (likelihood of high-degree nodes connecting to other high-degree degree nodes and low-degree nodes to other low-degree nodes), transitivity (if two nodes are connected, the probability that their adjacent nodes are also connected), edge count, density, mean degree, mean weighted degree (and adjusted), average shortest path length, and average weighted shortest path length (and adjusted). Node-level metrics comprised degree, closeness centrality, eigenvector centrality, PageRank centrality^36^, as well as their weighted counterparts, for assessing host species centrality.

We controlled for sampling effort through linear regression of edge weights, where the number of shared microbes between host pairs was predicted by the lesser sampling effort within each pair. Statistical significance of regression coefficients was explicitly tested to confirm positive associations between sampling effort and observed microbial sharing (Table S1). Original edge weights were then replaced with additively rescaled regression residuals constrained to positive values. All network metrics were subsequently recalculated using these adjusted weights^37,38^. Statistical significance of network parameters was assessed against 10,000 randomized null models generated through edge-weight permutations to determine whether the observed networks differed significantly from random networks^39^. To assess differences in the normalized adjusted weighted centrality metrics across the three orders, we again used PGLS, employing the same approach as for microbial richness. To quantify centrality, principal component analyses (PCA) were applied to the four adjusted weighted centrality metrics. The first principal component (PC1), explaining > 94% of total variance (Table S2), was adopted as a composite centrality index representing microbial transmission capacity^37^. Given the phylogenetic conservatism in traits regulating pathogen exposure, susceptibility, and suitability^40,41^, we quantified phylogenetic signal for microbial transmission capacity (represented by PC1 scores derived from centrality metrics) using Pagel’s λ^32–34^.

Microbe-sharing communities were identified using the Louvain algorithm from the *igraph* package^42–44^. The analysis was performed with edge weights set to NULL (as weights were inherently incorporated during network construction) and a default resolution parameter of 1 to balance community sizes. Final community assignments for all nodes were extracted and visualized using Cytoscape (v3.10.2).

### Ecological analyses

We next determined the ecological factors influencing microbial richness and transmission across the three host orders studied. Data on ecological factors were compiled from public databases, including life-history traits (body mass, litter size, gestation time, maximum longevity, and sexual maturity age), distribution features (habitat diversity and geographic range area), dietary diversity, urban adaptation status, minimum human population density (MHD) within distribution areas, and migratory behavior (for bats only). Body mass and gestation time data were sourced from the EltonTraits^45^ and AnAge databases^46^, respectively. Data on maximum longevity, sexual maturity age, geographic range area, and MHD were obtained from the PanTHERIA database^47^. Litter size data was sourced from the AnAge and Amniotes database^46,48^. Dietary diversity was quantified using the Shannon diversity index based on proportional representations across ten food categories in EltonTraits, calculated using the *vegan* package^29,45^. Habitat diversity was defined as the number of first-level habitat categories utilized by a species recorded in the International Union for the Conservation of Nature (IUCN) Red List database (https://www.iucnredlist.org/)^49^. Urban-adaptation status was determined using a published database on long-term urban adaptation in mammals^50^. Species classified as "dweller" or "visitor" were considered "urban-adapted" (1), while those not included in the urban-adapted dataset were coded as "non-urban" (2). Migratory behavior among bats was categorized as sedentary/short-distance, regional, or long-distance migratory^7^. Missing trait data were imputed using random forest approaches^51,52^.

PGLS models were applied to examine the effect of the above host traits on microbial richness and transmission^53,54^. Candidate models were constructed by including as many main effects as possible (i.e., body mass, litter size, gestation time, maximum longevity, sexual maturity age, habitat diversity, geographic range area, dietary diversity, urban adaptation status, MHD, sampling effort, and bat-specific migratory behavior), removing collinear predictor variables (i.e., Spearman’s ρ > 0.65; Fig. S1–Fig. S4; or variance inflation factor [VIF] > 5). The number of predictor variables was limited to ensure that each estimated coefficient had at least 10 observations. Candidate models were compared using Akaike information criterion (AIC), derived Akaike weights, and averaged coefficients when the ΔAIC of multiple models was less than two^55^. Model averaging was performed using the *MuMIn* package^54,56^. We constructed PGLS models for the pooled host orders and for each host order individually to quantify the effects of host traits on microbial metrics.

### Microbial transmission risk map

Global microbial transmission risk was comprehensively assessed by integrating spatially explicit population abundance data of host species with centrality metrics of their microbe-sharing network. Raster layers of species distributions and population abundance for both baseline and future scenarios were utilized^57–59^. The 2015 baseline data represented habitat suitability under 2000–2030 averaged climate conditions. Future projections modeled conditions for 2035 under two shared socioeconomic pathways (SSPs): SSP1 (sustainability development pathway, low climate stress) and SSP3 (regional rivalry pathway, high climate exposure), based on the 2020–2050 averaged climate conditions^60^.

Muicrobial transmission risk (*T_grid_*) was quantified per grid cell using the formula:

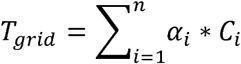

where *α_i_* is the population abundance of species *i* for both baseline and future scenarios^57–59^, *C_i_* denotes the standardized network centrality score (PC1) derived from microbe-sharing network topology, and *n* represents the number of host species within the grid cell. All analyses utilized a unified 0.5° geographic grid (WGS84, 55 km at the equator). Transmission high-risk hotspot areas were defined as grid cells with *T_grid_* values exceeding the 90^th^ percentile of global distribution. Emerging risk areas were grid cells where *T_grid_* increased by >20% from 2015 to 2035. Total and human-associated microbial transmission risk were quantified across the pooled host orders and for each host order. Geographic visualization was generated using national boundary vector data extracted via the *rnaturalearth* package and rendered with *ggplot2*^34^. To evaluate associations between predicted microbial transmission risk and zoonotic emerging infectious diseases (EID) risk, we fit a generalized linear mixed model (GLMM) with a beta distribution using the *glmmTMB* package^61^. The dependent variable, zoonotic EID risk (quantified as the probability of an EID event occurring within a grid cell, adjusted for reporting effort and human population), was derived from observed wildlife disease outbreaks between 1970 and 2016^62^. The independent variable was the predicted *T_grid_* values for human-associated microbes derived from the combined target host orders in 2015. Continent was included as a random effect to account for inherent regional heterogeneity in EID risk.

## Results

### Insectivores host comparable microbial richness to rodents and bats

We compared microbial diversity across insectivores, rodents, and bats. NMDS and PERMANOVA analyses revealed significant differences in both total (stress value = 0.161, *R*^2^ = 0.014, F = 9.29, df = 1341) and human-associated microbial compositions (stress value = 0.123, *R*^2^ = 0.022, F = 10.486, df = 915) across three host orders (all *P* < 0.001; Fig. 1b and Fig. 1c). To assess potential sampling biases, we used rarefaction curves, which showed accumulation of total microbial, viral, and bacterial richness with increasing sampling effort at the order levels (Fig. S5). Next, using PGLS, we found low phylogenetic dependence in microbial richness (λ range = 0–0.043; Table S3), while increased sampling effort universally enhanced richness across all host orders (PGLS: all *P* < 0.001; Table S3).

Following adjustment for sampling effort, our analyses revealed broad equivalence in microbial richness among insectivores, rodents, and bats. For human-associated microbes, insectivores exhibited richness comparable to both rodents (PGLS: β = −0.184, *P* = 0.095) and bats (β = −0.019, *P* = 0.865; Fig. 1e). This pattern extended to human-associated viral and bacterial subgroups, with insectivores showing richness comparable to both rodents (viruses: β = −0.078, *P* = 0.699; bacteria: β = −0.276, *P* = 0.067) and bats (β = 0.296, *P* = 0.122; β = −0.128, *P* = 0.434; Table S3). When examining total microbes, insectivores exhibited comparable richness to bats (β = 0.142, *P* = 0.108; Fig. 1d) and higher richness than rodents (β = −0.171, *P* = 0.047). Total viral and bacterial richness further supported this pattern of equivalence, revealing no significant differences among the three host orders (*P* > 0.05; Table S3).

### Higher cross-order and intra-order microbial transmission efficiency in insectivores

We constructed integrated microbe-sharing networks by combining data from insectivores, rodents, and bats, revealing distinct topological structures across host orders. Notably, insectivores showed a higher proportion of cross-order transmitted microbes (27.22%) than rodents (9.30%, χ^2^ = 228.54, *P* < 0.001) or bats (6.47%, χ^2^ = 370.53, *P* < 0.001; Fig. 2b and Fig. S6a). For human-associated microbes, insectivores maintained an even greater cross-order transmission proportion (75.90%) compared to bats (58.02%, χ^2^ = 86.74, *P* < 0.001) and rodents (37.56%, χ^2^ = 15.71, *P* < 0.001; Fig. 2a and Fig. 2b). Specifically, insectivores disproportionately shared microbes with rodents, accounting for 55.49% of their total microbial connections. Notably, this proportion increased to 59.38% when only considering human-associated microbes, indicating enhanced linkage specifically in microbes of zoonotic concern. Rodents primarily shared microbes with bats (46.69%) in the total microbe-sharing network, but displayed stronger intra-order connectivity for human-associated microbes, accounting for 45.70% of their connections. Bats, by contrast, maintained the highest intra-order connectivity in both total (50.95%) and human-associated microbe-sharing networks (54.09%), showing higher intra-order connectivity than cross-order transmission.

**Figure 2.**
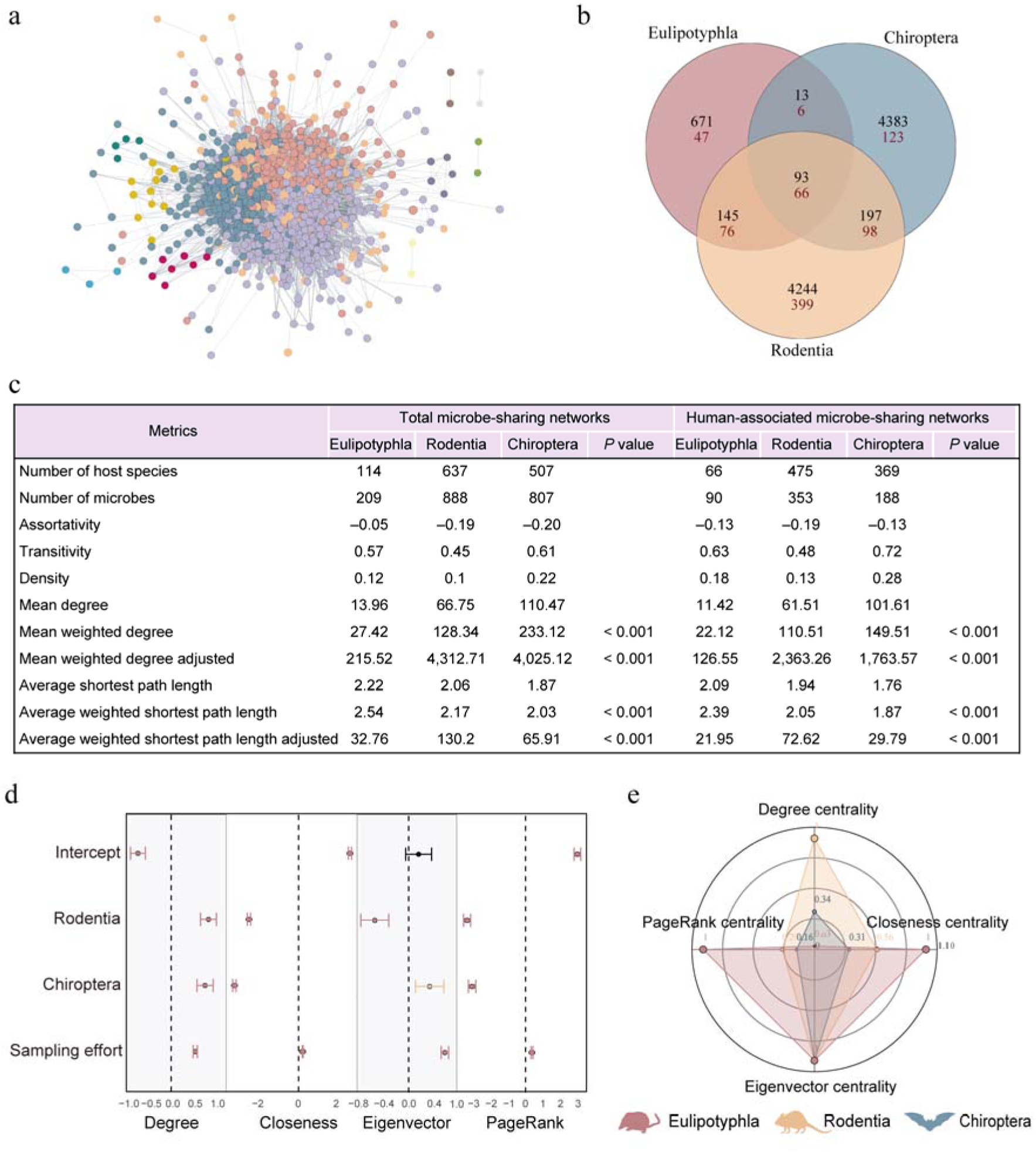
Microbe-sharing networks among insectivores, rodents, and bats. a) Human-associated microbe-sharing network for the pooled host orders (insectivores, rodents, and bats), with colors representing distinct community modules. b) Venn diagrams show the number of microbes shared among insectivores, rodents, and bats. Black numerals denote the number of shared total microbial species, while red numerals indicate the number of shared human-associated microbial species. c) Statistical metrics and associated *P*-values for the microbe-sharing network. *P*-values indicate the significance of difference between insectivores, rodents, and bats networks based on random permutations of the networks. *P* < 0.001 indicates significant differences between orders (pairwise tests). d) The coefficients estimates and 95%CIs for adjusted weighted centrality metrics of human-associated microbe-sharing networks derived from the PGLS models (*n* = 774). Circles represent coefficient estimates, and lines indicate 95%CIs. Statistical significance: red, *P* ≤ 0.001; yellow, *P* ≤ 0.01; blue, *P* ≤ 0.05; black, *P* > 0.05. e) Radar plots illustrating standardized adjusted weighted centrality metrics for central species in the human-associated microbe-sharing networks.

We also constructed order-specific networks to comparatively analyze microbial sharing patterns across these three orders, finding higher transmission efficiency within insectivores.

In human-associated microbe-sharing networks, insectivores showed higher network assortativity (−0.131) than rodents (−0.19) and bats (−0.130; Fig. 2c). Insectivores exhibited intermediate network density (0.18) and transitivity (0.63) between rodents (density: 0.13; transitivity: 0.48) and bats (0.28; 0.72; Fig. 2c). Notably, following adjustment of sampling effort, insectivores showed significantly shorter mean weighted shortest path lengths (21.95) than rodents (72.62) and bats (29.79; Fig. 2c). We found similar patterns in the total microbe-sharing networks (see Supplementary Information).

Furthermore, centrality analyses identified central species acting as hubs within the host order-specific microbe-sharing networks. PGLS models comparing four adjusted weighted centrality metrics revealed a weak-to-moderate phylogenetic signal (λ range = 0.147–0.694; Table S4). In both total and human-associated microbe-sharing networks, insectivore species consistently exhibited significantly higher normalized adjusted weighted closeness and PR centrality scores than those of rodents or bats (PGLS: all *P* < 0.001; Fig. 2d and Fig. S6c). Examining specific central host species, the European shrew (*Sorex araneus*) in the total network exhibited superior adjusted weighted closeness, eigenvector, and PR centrality compared to the black rat (*Rattus rattus*) and common vampire bat (*Desmodus rotundus*; Fig. S6b). Concordantly, within human-associated networks, the European shrew consistently emerged as the most central species, showing significantly higher adjusted weighted closeness and PR centrality than the house mouse (*Mus musculus*) and Leschenault’s rousette (*Rousettus leschenaultii*; Fig. 2e).

### Ecological drivers of microbial richness and transmission

To further assess the effects of ecological factors on microbial richness and transmission across and within the three host orders studied, we again used PGLS models. Our analyses revealed limited phylogenetic influence on microbial richness and transmission. Across all three orders, we found low phylogenetic signal for total (λ = 0.085) and human-associated microbial transmission capacity (λ = 0.067). Additionally, order-specific models revealed that insectivores showed weak phylogenetic signal in human-associated microbial richness (λ = 0.029), while rodents exhibited weak phylogenetic signals in both total (λ = 0.187) and human-associated microbial transmission capacity (λ = 0.161). Bats displayed weak phylogenetic signal only for total microbial richness (λ = 0.004).

For data pooled across host orders, urban-adaptation status, maximum longevity, and geographic range area all were positively correlated with human-associated microbial richness and transmission capacity (Fig. 3a, Fig. 3b, and Table S5). Additionally, habitat diversity (β = 0.1, *P* < 0.001) and sampling effort (β = 0.85, *P* < 0.001) had a positive effect on human-associated microbial richness, while body mass had a negative effect (β = −0.56, *P* < 0.001). For human-associated microbial transmission capacity, litter size had a positive effect (β = 0.22, *P* < 0.001) while sexual maturity age had a negative effect (β = −0.19, *P* < 0.001).

**Figure 3.**
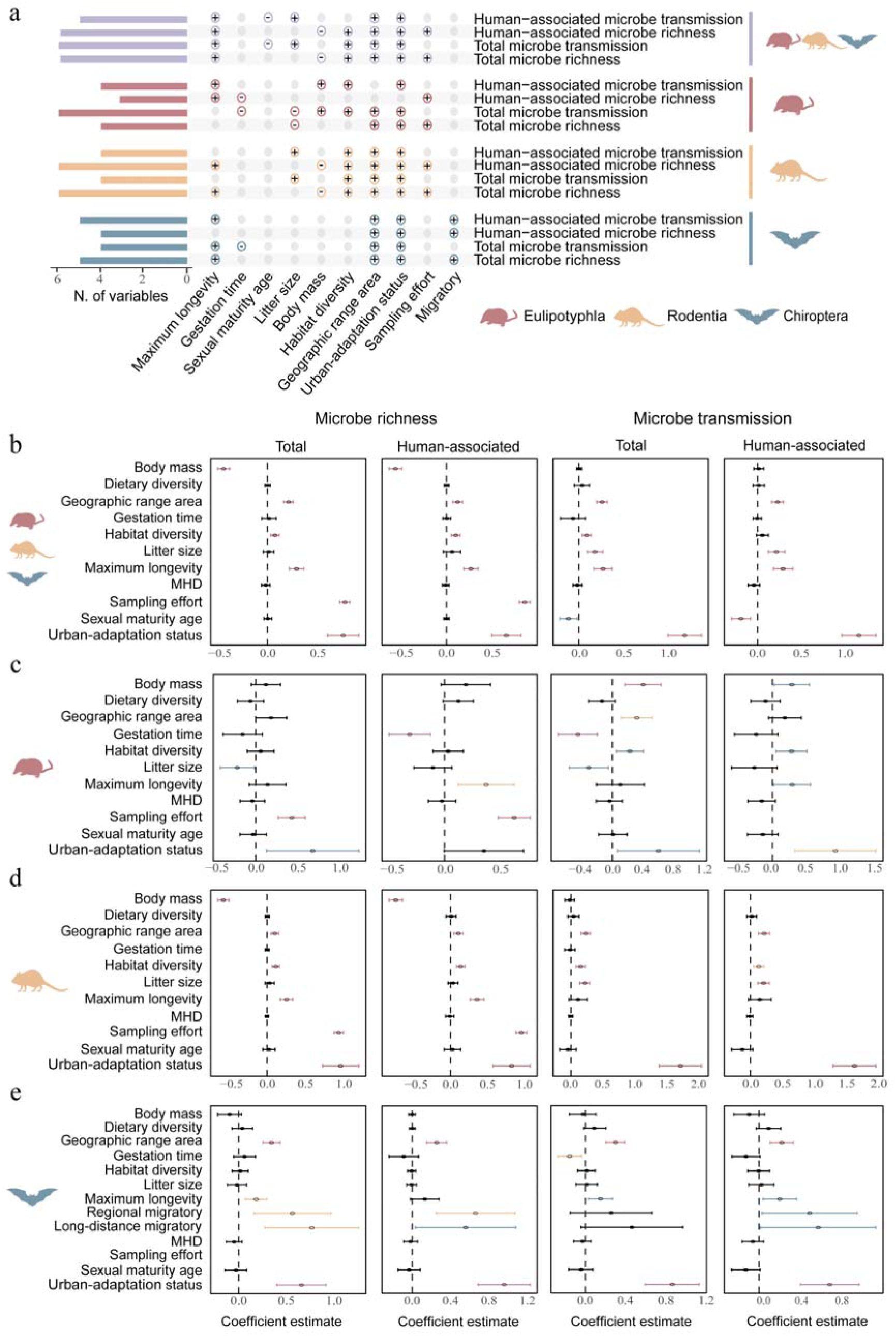
Influence of ecological traits on microbial richness and transmission in insectivores, rodents, and bats. a) Summary of PGLS analyses. Bar plots (left) indicate the number of significant ecological traits associated with microbial richness or transmission. Dot plots (middle) identify influential ecological traits, with the symbols denoting a positive (+) or negative (−) effect. Colors represent distinct host orders. The coefficients estimates and 95%CIs for microbial richness and transmission derived from the best PGLS models in b) the pooled host orders, c) insectivores, d) rodents, and e) bats. Circles represent coefficient estimates, and lines indicate 95%CIs. Statistical significance: red, *P* ≤ 0.001; yellow, *P* ≤ 0.01; blue, *P* ≤ 0.05; black, *P* > 0.05. MDH represents minimum human population density within distribution areas.

Order-specific analyses showed that both geographic range area and urban-adaptation status were generally positively correlated with human-associated microbial metrics (e.g., the richness and transmission capacity; Fig. 3a and Table S6–Table S8). Habitat diversity was also generally positively associated with the human-associated microbial metrics in both insectivores and rodents (Fig. 3c and Fig. 3d) but not bats (Fig. 3e). For life-history traits, longevity was generally positively correlated with the human-associated microbial metrics in all three orders. Gestation time showed negative relationships with human-associated microbial richness in insectivores as well as with total microbe transmission capacity in both insectivores and bats. In contrast, body mass and litter size showed order-specific effects (Fig. 3a, Fig. 3c and Fig. 3d). Increased body mass enhanced human-associated microbial transmission capacity in insectivores (β = 0.28, *P* = 0.031) but suppressed human-associated microbial richness in rodents (β = −0.74, *P* < 0.001). Similarly, litter size positively correlated with human-associated microbial transmission capacity in rodents (β = 0.21, *P* < 0.001). In addition, migration behavior was uniquely associated with human-associated microbial metrics in bats (Fig. 3e). We observed similar patterns of ecological drivers for microbial richness and transmission across both total and human-associated microbes (see Supplementary Information). Best-fit PGLS models are detailed in Tables S9–S12.

### Expanding hotspots of zoonotic risk

We conducted spatially explicit analyses to quantify human-associated microbial transmission risks across insectivores, rodents, and bats for the 2015 baseline (representing 2000–2030 averaged climate conditions) and 2035 projections (2020–2050 averaged climate conditions) under SSP1 and SSP3 scenarios. The 2015 baseline revealed order-specific risk patterns. Insectivore-associated, high-risk hotspots primarily occurred in Eurasia (Eastern/Southern Europe and East/Southeast Asia; Fig. 4b). Rodent-associated hotspots were concentrated in Eurasia (Southern/Central Europe and East/South/Southeast Asia) and North Africa (Fig. 4c). Bat-associated hotspots were prominent in tropical regions, including northern South America, Central Africa, and coastal Southeast Asia (Fig. 4d). Overall, hotspots for the combined target host orders exhibited spatial convergence with the rodent-associated risk patterns (Fig. 4a).

**Figure 4.**
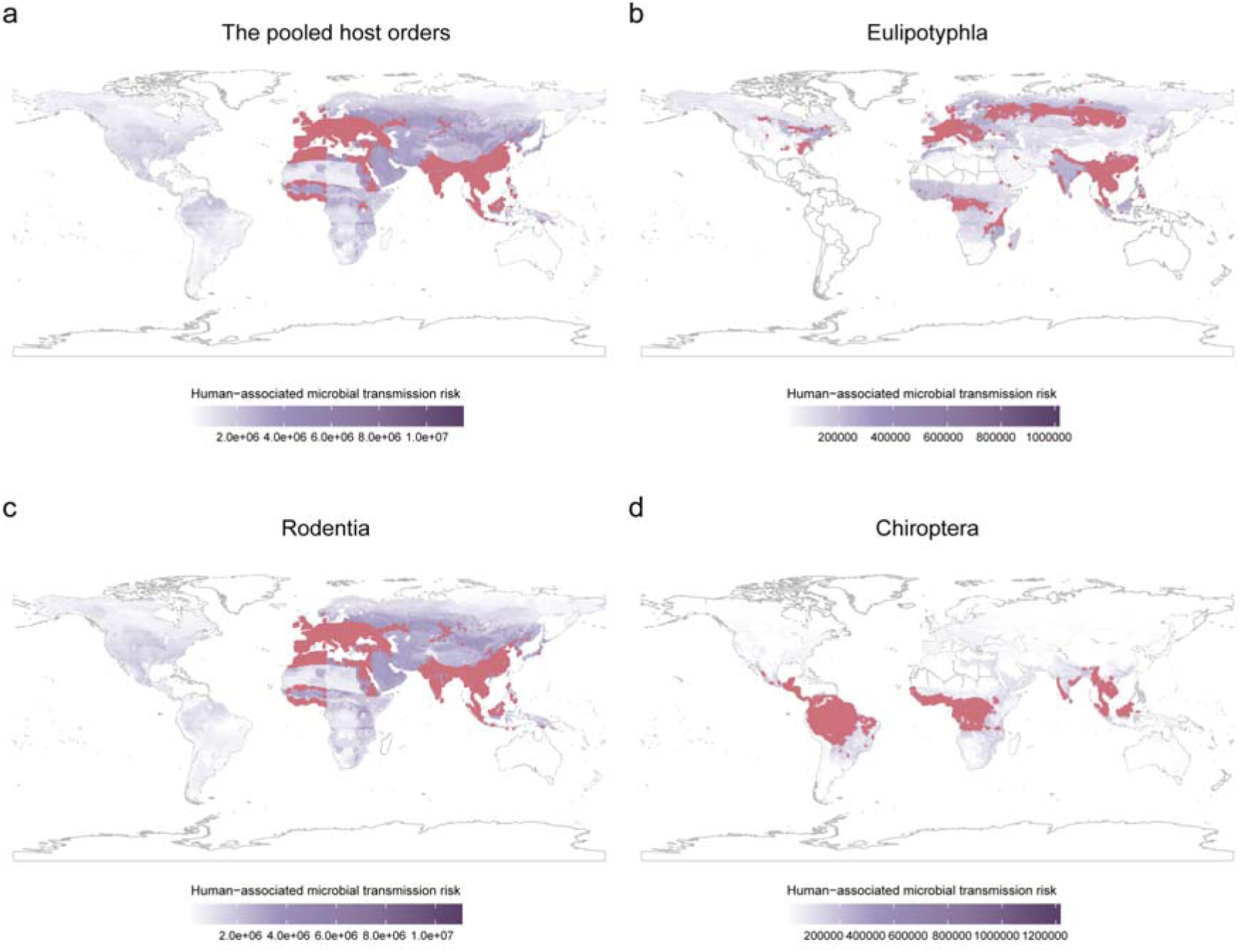
Global patterns of human-associated microbial transmission risk in insectivores, rodents, and bats in 2015. Predicted spatial distribution of hotspots for human-associated microbial transmission risk in a) the pooled host orders, b) insectivores, c) rodents, and d) bats. Darker shading indicates higher transmission risk, and red areas represent high-risk hotspot regions.

Projected risk landscapes for 2035 under SSP1 revealed a substantial expansion of high-risk areas (Fig. 5, Fig. S7–Fig. S10). Notably, insectivores emerged as the predominant transmission host in new high-latitude hotspots, including parts of the United States, Canada, and Russia, surpassing the relative risk contribution of rodents and bats (Fig. 5e). In addition to the persistence of core 2015 hotspots, this order also demonstrated expansive emerging risk across temperate regions of East Asia, North America, and Europe (Fig. 5c). In contrast, rodent-associated emerging risk areas developed primarily in southwestern North America, South America, Central/West Africa, and Australia (Fig. S9). Bat-associated emerging risk areas were concentrated mainly in Eastern/Southern Africa, East Asia, and Europe (Fig. S10). Overall, emerging risk areas for the combined target host orders showed spatial patterns similar to rodents (Fig. 5a). SSP3 projections indicated less spatial expansion of risk areas than SSP1 (Fig. 5, Fig. S8–Fig. S10). The spatial distributions of transmission risk from total and human-associated microbes were highly similar (Fig. S7–Fig. S10).

**Figure 5.**
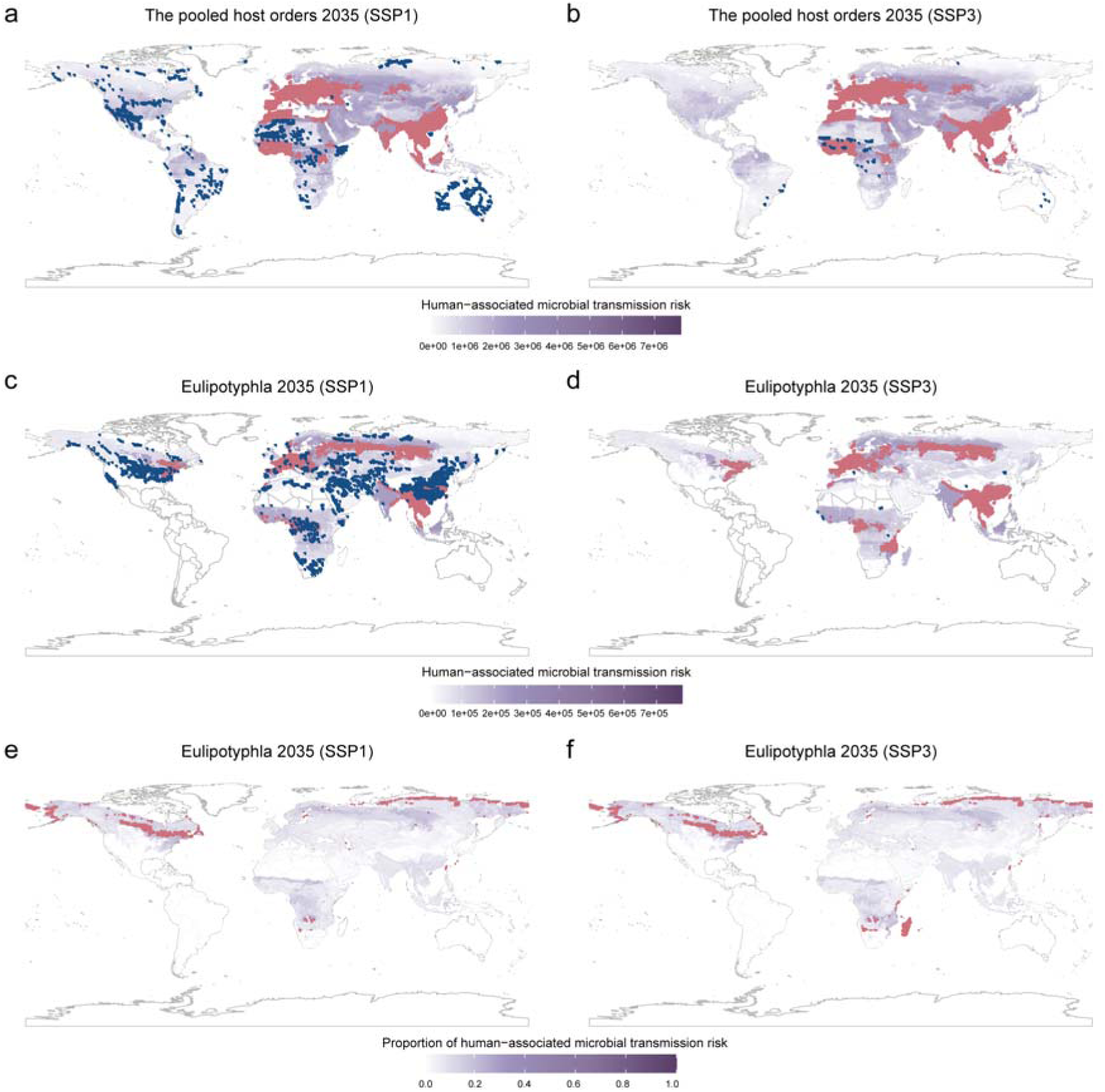
Projected global patterns of human-associated microbial transmission risk in insectivores, rodents, and bats under 2035 climate change scenarios. Predicted spatial distribution of hotspots for human-associated microbial transmission risk in the pooled host orders under a) SSP1 and b) SSP3. Predicted spatial distribution of hotspots for human-associated microbial transmission risk in insectivores under c) SSP1 and d) SSP3. In panels a)-d), darker shading indicates higher transmission risk; red areas represent high-risk hotspot regions, while blue highlights areas with >20% increased transmission risk between 2015 and 2035, suggesting emergent risk zones. The proportion of human-associated microbial transmission risk attributable to insectivores among the pooled host orders under e) SSP1 and f) SSP3. Regions in red (proportion >0.34) denote areas where insectivores are predicted to be the dominant source relative to rodents and bats.

Finally, using a global database of historical EID events (1970–2016), we analyzed associations between predicted human-associated microbial transmission risk in 2015 and global EID risk. Our GLMM with continent as a random factor explained 95.36% of global EID risk variation (conditional *R*^2^), with 20.32% explained by microbial transmission risk (marginal *R*²; β = 0.316, *P* = 0.028). This provides strong empirical validation that areas with higher predicted transmission risk in our target host orders were significantly more likely to have experienced emerging disease outbreaks in the recent past. Notably, predictive performance varied by continent, with Europe (44.3%) and North America (39.8%) showing the strongest associations, while Oceania (7.9%) and Asia (0.149%) exhibited minimal explanatory power.

## Discussion

### Insectivores as equally critical microbe reservoirs

In this study, we took a macroecological approach to investigate microbial hosting and transmission dynamics across insectivores, rodents, and bats. Counter to the prevailing paradigm favoring rodents and bats as special reservoirs of zoonotic pathogens^11^, our analyses demonstrate that insectivores host comparable richness of microbes to rodents and bats. Moreover, insectivores exhibited superior efficiency not only in intra-order transmission, but also in cross-order transmission, relative to bats and rodents. These findings establish insectivores as pivotal yet neglected players in disease ecology, compelling their systematic incorporation into global zoonotic risk assessments.

Our findings align with emerging evidence identifying insectivores as significant reservoir hosts for multiple pathogens ^63–65^. Hedgehogs, for example, have been recognized as amplifying hosts for SFTSV ^21,66,67^, which the World Health Organization has included on its list of priority pathogens with pandemic potential in both 2017 and 2024. Hedgehogs may also constitute a reservoir for methicillin-resistant *Staphylococcus aureus* carrying *mecC*^68,69^. Shrews have been considered as natural reservoirs of the Langya henipavirus, which is a close relative of the Hendra and Nipah viruses^22^. Certain insectivore species (e.g., Smith’s shrew [*Chodsigoa smithii*]) host more viruses than some species of rodents and bats^3^. Furthermore, our recent work demonstrated that insectivores, specifically shrews and hedgehogs, contribute significantly to the global virus sharing networks among mammal species, hosting many cross-order transmitted viruses^8^.

### Ecological drivers of microbial richness and transmission

We explored and compared the ecological factors driving microbe richness and transmission capacity across these three host orders, identifying both shared and order-specific drivers. For example, most habitat-related factors showed positive correlations with microbe richness and transmission capacity in all orders, except habitat diversity, which showed no relationship in bats. These positive effects may generally reflect increased microbe exposure, a key process in host–microbe interactions^70,71^. Species with larger geographic ranges, greater habitat diversity, and urban adaptation also experienced higher microbe contact, which could likewise indicate greater within- and between-species transmission opportunities, including spillback from humans^72^. Additionally, longevity generally correlated positively with microbe richness and transmission capacity across all groups, likely because long-lived species have greater cumulative exposure to microbes^73,74^.

Besides the shared drivers, we also identified several order-specific drivers for species’ microbial richness and transmission capacity. For rodents, we found that species with smaller body mass and larger litter sizes, and therefore a faster pace of life, showed higher microbial richness and transmission capacity. These results are thereby consistent with life-history theory, which posits that such fast-lived species typically invest less in immune defense than in reproduction and are thus more susceptible to microbes^75,76^. However, we found that the effects of body mass and litter size in insectivores were opposite to those in rodents. Such opposing patterns in insectivores might be explained by exposure processes. Within insectivores, small-bodied species often exhibit strict insectivory due to their high metabolic rates^77^. In contrast, larger-bodied species (e.g., hedgehogs^78,79^), owing to their greater absolute energy requirements, may have larger home ranges^80,81^ and exhibit opportunistic omnivory, which could expose them to a greater diversity of microbes. We also found that migratory bat species host more human-associated microbes than non-migratory bat species, which could likewise reflect such species sampling a greater diversity of habitats and, as such, being exposed to more microbes^82^.

### Implications of expanding transmission risks for public health preparedness

Our spatial projections based on climate change scenarios revealed a concerning expansion of insectivore-linked high-risk zones for microbial transmission by 2035. Under SSP1 climate scenarios, insectivore-driven spillover risk will increase across temperate regions of East Asia, North America, and Europe. Notably, insectivores are forecasted to surpass rodents and bats and emerge as the predominant transmission hosts in new high-latitude hotspots, including parts of the US, Canada, and Russia, likely driven by climate advantages in their range, abundance, or vector associations. The European shrew, identified as the most central species in microbial transmission networks, may amplify spillover threats and potentially trigger regional public health crises. Our data underscores how climate change fundamentally reconfigures host–microbe networks, elevating insectivores from neglected actors to central drivers of emerging spillover risk, demanding a commensurate shift in surveillance paradigms.

Our findings establish insectivores as critical hosts of underestimated microbial diversity with significant zoonotic potential, compelling their prioritization in proactive surveillance systems for emerging infectious diseases. We propose the immediate incorporation of insectivores into existing surveillance programs, with targeted focus on high-risk species such as the European shrew and on geographic hotspots to comprehensively map pathogen diversity and quantify zoonotic risk. Concurrently, integrating multi-omics approaches could reveal how insectivores’ biological traits contribute to their maintenance and transmission of diverse pathogens to advance mechanistic insights into zoonotic spillover^83^. Furthermore, implementing One Health sentinel networks that synergize environmental surveillance, point-of-care animal testing, and human digital epidemiology would establish early-warning systems across high-risk interfaces, translating ecological insights into public health intervention protocols^84^. Public health agencies in areas with high insectivore biodiversity should prioritize community education about safe handling of these animals. These collective actions will bridge the critical preparedness gaps exposed by neglected insectivores.

### Limitations

We applied rigorous measures to mitigate the impact of inherent data biases on our major findings. Our rarefaction curves and PGLS demonstrated that increased sampling effort enhanced microbial richness in insectivores, rodents, and bats. Independent analyses for human-associated microbes, viruses, and bacteria were conducted to address group-specific bias. Nevertheless, we acknowledge that inherent data biases, including temporal and geographic sampling gaps, may skew microbial richness and cross-species transmission comparisons. Additionally, findings on microbe-sharing dynamics and zoonotic potential warrant cautious interpretation, particularly as our network-based framework cannot conclusively exclude alternative transmission scenarios. Our analyses did not distinguish non-vector from vector transmission, although the latter represents an important transmission pathway for certain pathogens in wild small mammals. While our analysis delineates potential cross-species transmission risks, combined evidence from viral isolation and genomic sequencing, phylogenetic analysis, epidemiological contact tracing, and serological profiling are needed to validate definitive transmission routes.

## Conclusions

In summary, our findings demonstrate that insectivores pose a substantially greater zoonotic risk than previously acknowledged, evidenced by their comparable microbial diversity, higher transmission efficiency, and elevated spillover potential. Collectively, insectivores, rodents, and bats constitute essential zoonotic host orders, a role sustained by distinct ecological traits that promote pathogen maintenance and spillover. Consequently, proactive surveillance of these taxa is imperative for early warning of emerging zoonotic threats prior to spillover events.

## Acknowledgements

This study was supported by the National Natural Science Foundation of China (32470561, YFX, 32271605, ZYXH) and Taishan Scholars Project (tsqn202306003, YFX). DJB was supported by the National Science Foundation (DBI 2515340) and the Edward Mallinckrodt, Jr. Foundation.

## Competing interests

None declared.

## Notes

### Competing Interest Statement

The authors have declared no competing interest.

